# The complexity of keratocyte migration in salmon explant cultures: initial results and future prospects

**DOI:** 10.1101/2022.06.01.494312

**Authors:** Ida S. Opstad, Deanna L. Wolfson, Balpreet S. Ahluwalia, Krishna Agarwal, Tore Seternes, Roy A. Dalmo

## Abstract

Intact skin is of uttermost importance for fish welfare. The fish skin provides an environmental barrier and protects against invading pathogens. However, both pathogens and physical insults cause skin wounds that are of major concern in modern fish farming. The behavior and interactions between keratocyte cells and sheets of cells are not well understood. The collective migration of keratocytes (skin epithelial cells) is of central importance for wound healing in fish. In this study, we aimed to elucidate the complex wound healing process in fish skin by studying in vitro cultures of these highly motile cells. Using explant cultures from farmed Atlantic salmon (*Salmo salar*) and differential interference contrast microscopy (DIC), we have captured the dynamics of sheets of cells from harvested fish scales and of individual cells interacting in the cell sheet vicinity. In addition to direct contact, the cells were observed to interact through long membrane tubes, turn, rotate, merge, and/or detach. Additionally, stationary cells and cells moving on top of the cell sheets were observed. Cell sheets approaching one another from different scales did not merge but dispersed.

## 1. Introduction

The fish skin provides a barrier to environmental cues and against invading pathogens. Some infectious pathogens and physical insults cause skin wounds [1], which have to be re-epithelized and remodeled quickly to sustain the barrier function. In fact, small wounds may be healed within a few hours [2], depending on the environmental temperature. Keratocytes cover the dermis which is composed of dense connective tissue, fibroblasts, pigment cells and blood capillaries [3] [4]. Intact skin is crucial for animals’ defence against pathogens. In fishing industry, salmon louse infestation is a huge concern for animal welfare. Non-medicinal treatments include physical delousing processes such as freshwater bathing, warm water dips, use of lasers to kill individual lice on the fish, mechanical removal of parasites by soft brushes and/or high-pressure pumps [5]. Such physical treatments are likely to affect the fish skin integrity as well as the activity of keratocytes, which are important for tissue remodeling. The migratory behavior of keratocytes (skin epithelial cells) is a key feature during wound repair. It has been shown that the speed of the fish moga (*Hypsophrys nicaraguensis*) keratocytes is approximately 15 μm/min (23 °C) in vitro [6]. This finding is similar to the previous results presented by [7] where the velocity of black tetra (*Gymnocorymbus ternetzi*) keratocytes was calculated to be 15 μm/min. There are numerous studies on fish keratocytes that have described the migratory features of individual cells [8] [9] [10]. However, the far more complex behavior of sheets of cells – as found in fish skin – is comparatively unexplored. Here, cells are seen to transition between *leader* and *follower* cell morphology. The leader cells at cell-sheet edges typically exhibit the characteristic keratocyte fan-shape and appear to determine the direction of the entire sheet of cells. The follower cells are within the cell sheet and follow the movement of the neighboring cells [11] [12].

Using explant cultures from farmed Atlantic salmon (*Salmo salar*) and differential interference contrast microscopy (DIC), we have observed both the advance of sheets of cells coming off the harvested fish scales and the interactions of individual cells detaching and recombining both with other cells and with the cell sheet. In this work, we show videos of Atlantic salmon keratocytes acquired at room temperature, and discuss their complex dynamics as observed in scale explant cultures.

## 2. Materials and Methods

### 2.1. Cell harvesting and cultivation

The imaged cells were primary epidermal skin cells (mainly keratocytes) obtained from Atlantic salmon scales. The scale harvest was conducted with disinfected tweezers on farmed salmon in air, killed with a blow to the head. This method is allowed according to the Norwegian Regulations for use of animals in experimentation (https://lovdata.no/dokument/SF/forskrift/2015-06-18-761#KAPITTEL_10) and complies with the corresponding EU legislation - Directive 2010/63/EU (http://data.europa.eu/eli/dir/2010/63/oj). The scales were placed on #1.5 glass-bottom dishes (734-2904, VWR) and dried on for about 1-2 minutes (to make the scales stick to the dish) before adding antibiotic buffer solution: Hank’s Balanced Salt Solution (HBSS, Corning, #21-023-CM) with 100 IU/mL Penicillin-Streptomycin (P/S, Sigma, P0781) and 1 μg/mL Amphotericin B (PanReac AppliChem, #A7009). The short dry period did not appear to damage the harvested cells. The cells were kept at 4°C in a watertight box, replacing the old medium with fresh every 2-3 days. Scales that loosened during cell culture and sample preparation were removed from the culture dish. Before imaging, loose cells and mucus were washed away by changing the media to a fresh HBSS antibiotic solution.

### 2.2. Image acquisition

Imaging was performed at room temperature using a DeltaVision Elite microscope (GE Healthcare) equipped with a DIC module, and an Olympus 10X, 0.4NA or 20X, 0.75NA objective. Unless otherwise stated, the DIC module with the 10X objective was used. The microscope focus together with the contrast of the DIC module were manually optimized to maximize the visibility of the lamellipodia of leading-edge keratocytes adherent to the glass-substrate. For long-duration time-lapses, the keratocytes were maintained in sharp contrast with help of the microscope’s inbuilt focus maintenance system. The exact acquisition parameters for all raw image data are available with the published dataset [13].

### 2.3. Image processing and analysis

All image processing (linear contrast adjustment, file conversion, insertion of scale bars, etc.) was conducted in Fiji [14]. The stitched images (combining multiple smaller field-of-views, files labeled STC) were generated using the Fiji plugin Grid/Collection Stitching [15].

## 3. Results

With the explant culture preparation described under Material and Methods, two different image acquisition schemes were conducted over 0-4 days after sample harvesting:

I. 2 min time-lapses (for up to about 24 hours) often consisting of multiple adjacent images together covering a large sample area (e.g. 3 × 3 mm^2^.) where cells (or sheets of cells) were starting to come of the scale (i.e., a beginning cell avalanche).
II. 5 s time-lapses (for up to 14 hours) covering a single field-of-view (either 668 × 668 μm^2^ for 10X lens, or 334 × 334 μm^2^ using a 20X lens) close to a scale edge.

Approach I enabled an overview of the scale avalanche (large sheet of keratocytes) dynamics over a relatively long time-frame and large field-of-view to be captured, while at the same time achieving sufficient image resolution and contrast to capture individual cell dynamics, including the very thin lamellipodia of the cell sheet leader cells, as demonstrated in Figure 1. Video 1 shows an overview of the full avalanche dynamics. Similarly, Video 2 shows an overview of a different sample, but here of two approaching avalanches from two different salmon scales (but from the same fish). Surprisingly, such avalanches were not found to merge but rather tended to disperse or avoid each other. Considering that cell avalanches *in vivo* are supposed to cover skin wounds, we expected to see the avalanches merging. The temperature or phototoxicity for imaging could be important contributors to this behavior, especially since we found multi-scale dishes (kept at 4°C protected from light) to eventually be fully covered by cells after enough days on a dish.

**Fig. 1.**
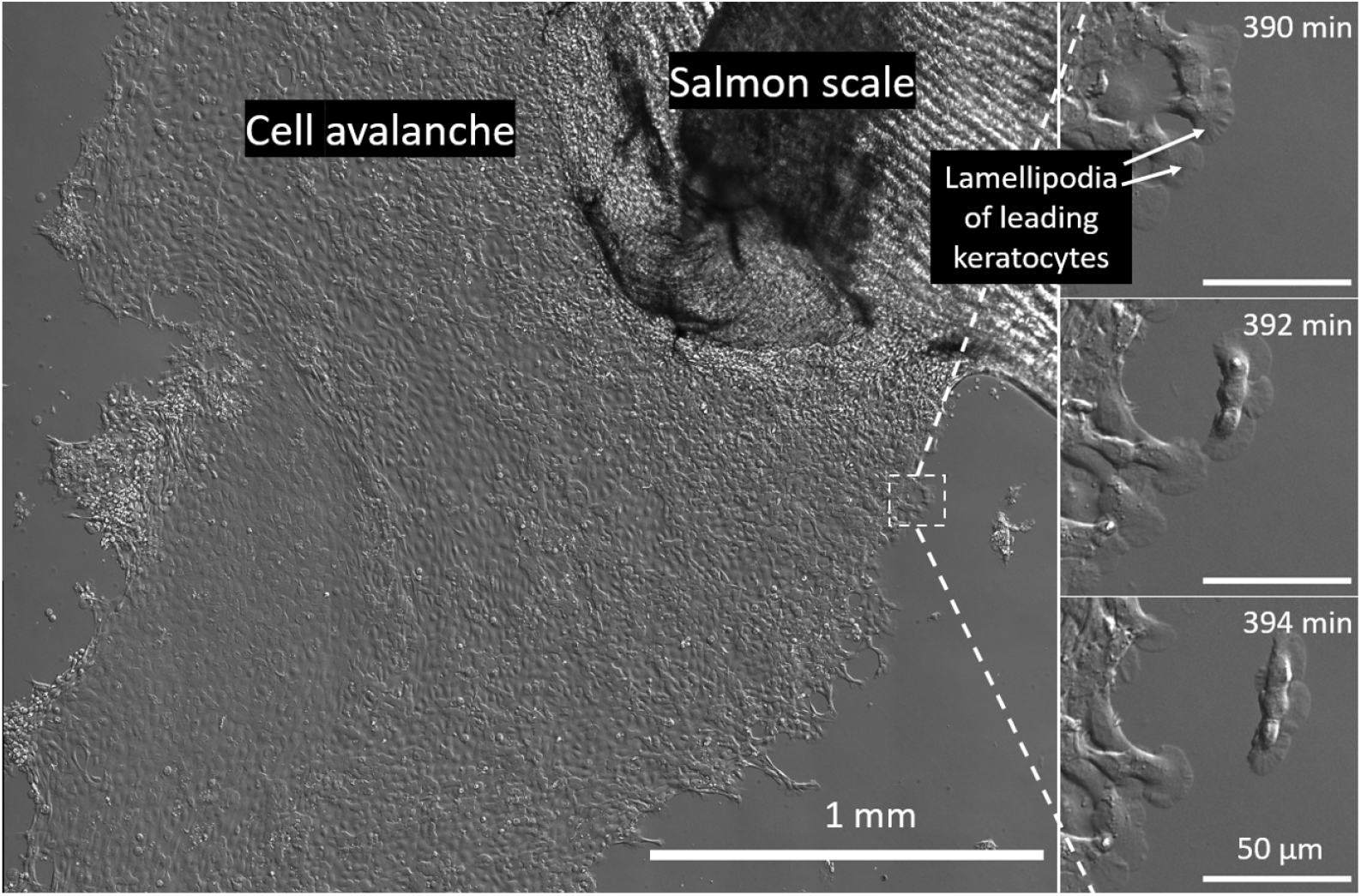
Live imaging of explant scale culture of Atlantic salmon 3 days after harvest. Epithelial cell avalanches (as seen coming off the scale in the main panel) are important in the wound healing mechanisms of fish skin. This approach enabled an overview of the explant culture cell avalanche to be captured at the same time as achieving sufficient resolution and contrast to investigate individual cell and lamellipodium dynamics (right panels). The left panel is a tiled view of 5 × 6 DIC images. The cell avalanche shown had already been imaged for 390 min (every 2 min) at room temperature. No nutrition was provided to the culture medium (antibiotic-supplemented HBSS).

With approach II, we were better able to study cell-cell interaction and details of their complex behaviour. The following section outlines some interesting observations regarding individual and inter-cell interactions following approach II.

### 3.1. Cell dynamics (5 s time-lapse)

Different from the unidirectional motion normally described following keratocyte analysis, our explant culture data show a broad range of different cell behavior and motion patterns, as outlined in Figure 2. Some changes in propagation direction are clearly induced by direct contact with other cells or membrane tubes (see Video 3), but also single cells without no apparent direct cell contact were found to change the direction of propagation, as can be seen from the time-lapse displayed in Figure 3 and Video 4, where a cell turns following apparent folding of its lamellipodium. Figure 4 and Video 5 display a rotating cell (or collection of cells), seemingly caused by a combination of keratocyte cells migrating in opposite directions.

**Fig. 2.**
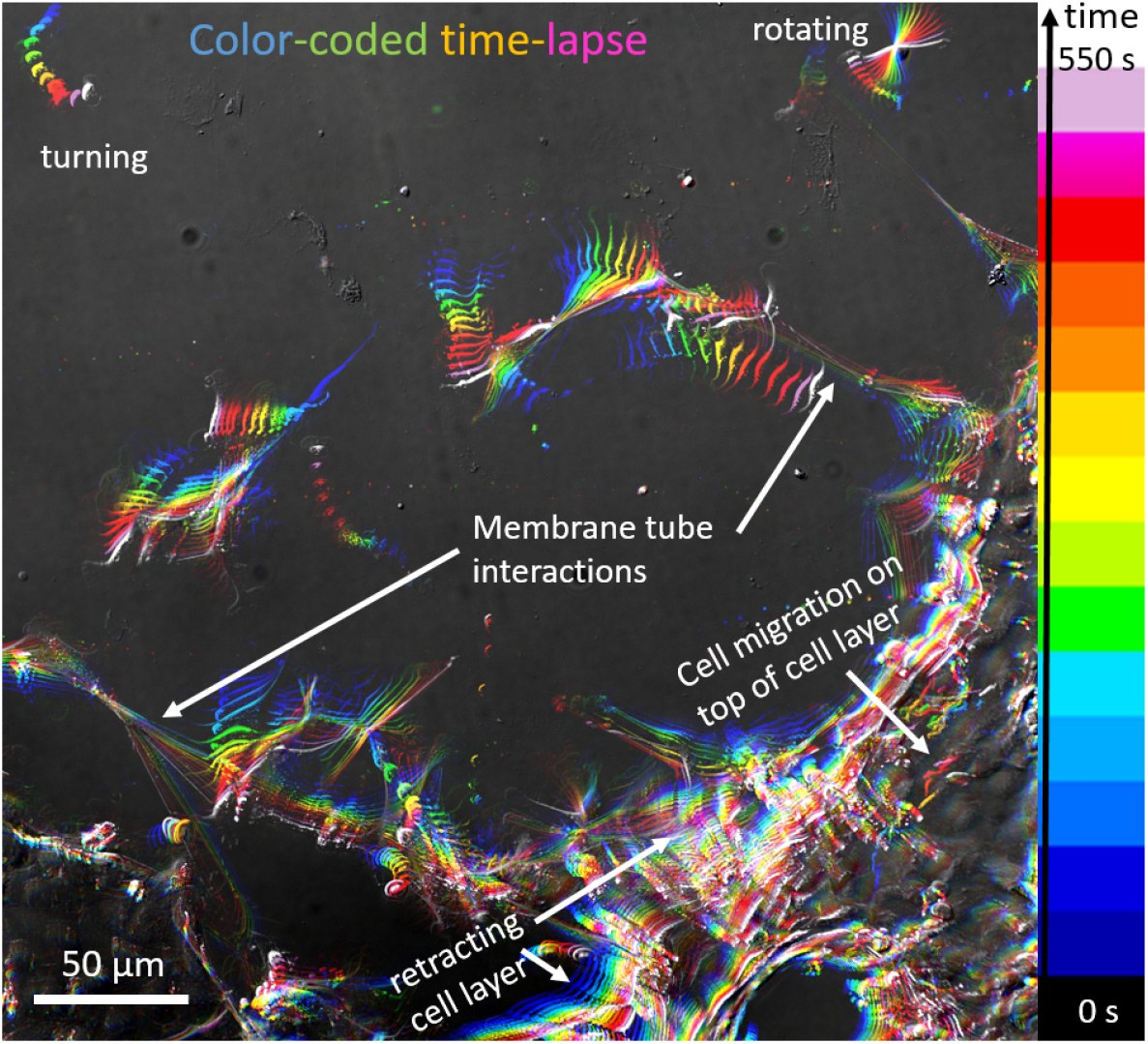
Color-coded time-lapse of keratocyte video: each color represents a different time-point according to the sequence indicated by the color bar (50 s between each color). Cells undergo several different dynamical patterns: cells are turning, rotating, interacting through membrane tubes, and cells are moving on top of the retracting cell layer. Scale bar: 50 μm. The figure was created with help of a Temporal-Color Coder script, provided by Kota Miura at the Centre for Molecular and Cellular Imaging, EMBL Heidelberg, Germany.

**Fig. 3.**
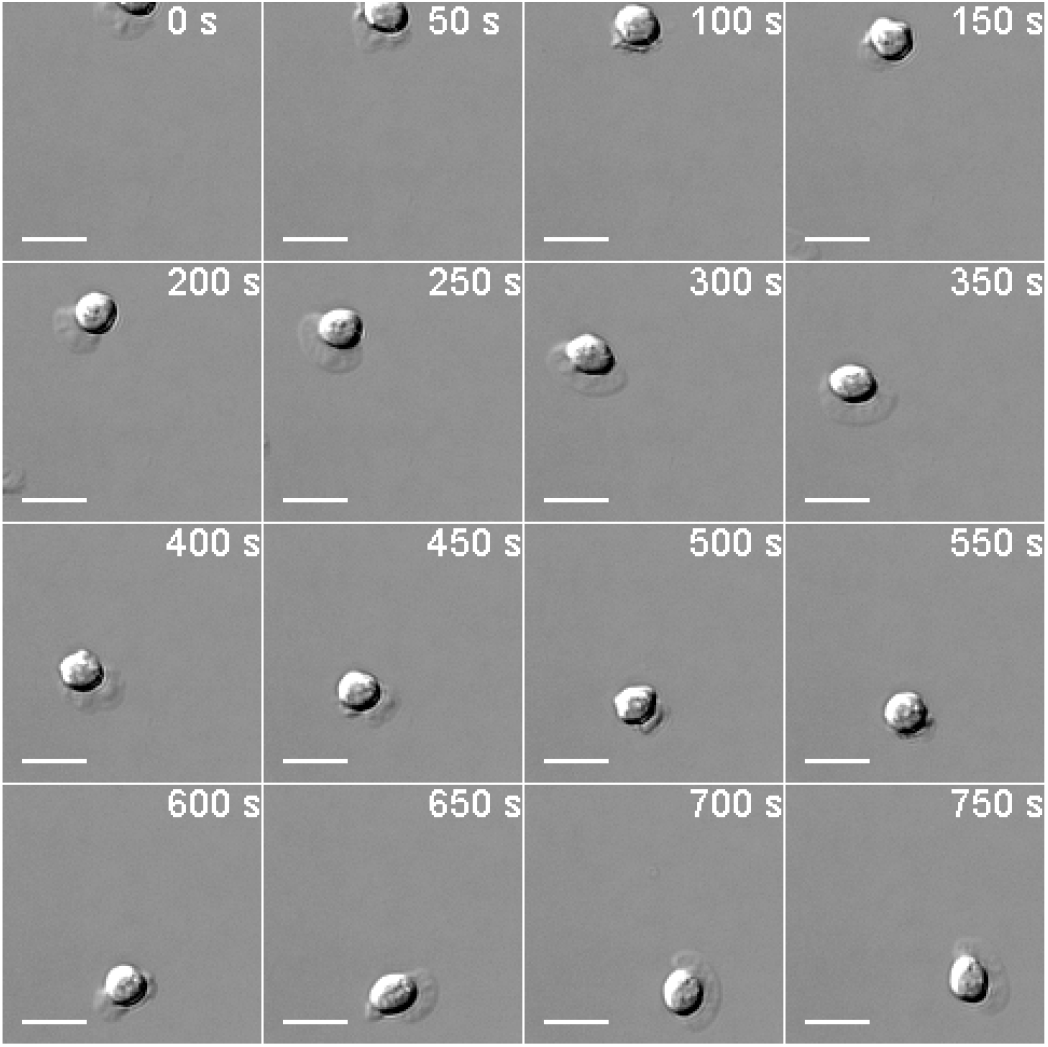
Time sequence of a cell balling up and turning. Scale bars: 20 μm.

**Fig. 4.**
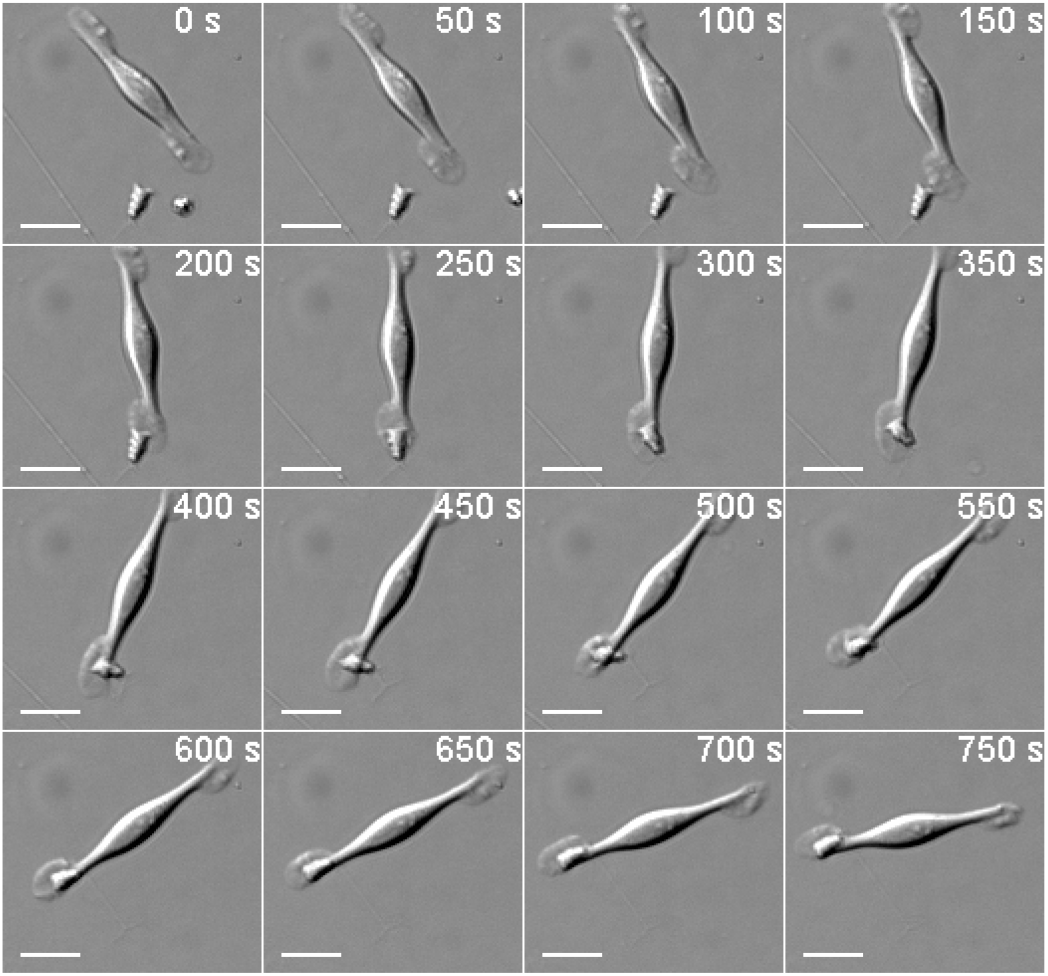
Time sequence of a rotating cell collecting and transporting along a large particle from 200 s – 750 s. Scale bars: 20 μm.

Interestingly, also cells moving independently on top of the keratocyte cell sheet were observed (Video 6 and indicated by yellow arrows in Figure 5). These cells appear different from keratocyte cells in migratory behavior and morphology, although they have not been further characterized beyond the visual observation.

**Fig. 5.**
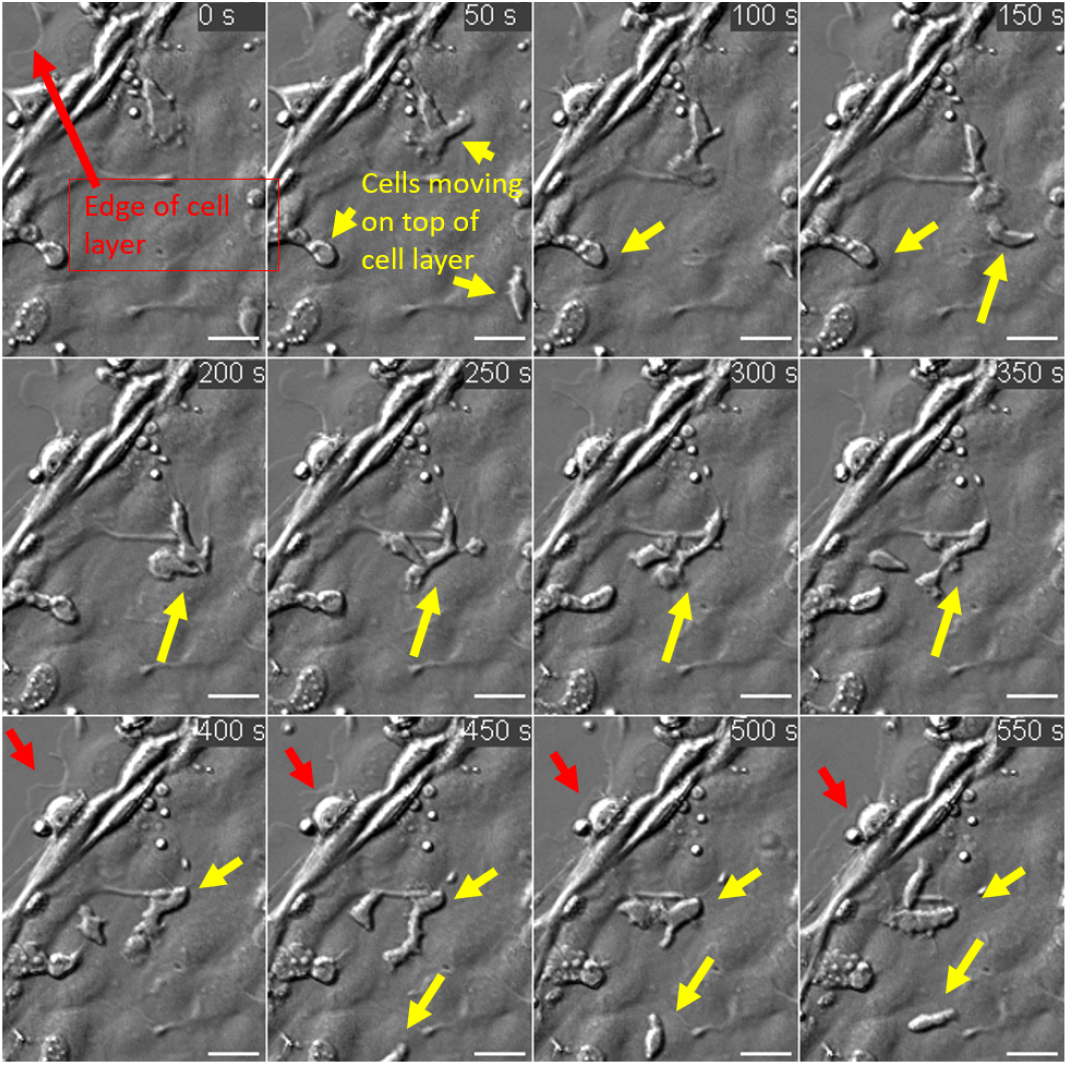
Time sequence of cells moving on top of a connected but dynamical cell layer. The red arrows point at the edge of the cell sheet, while the yellow arrows are indicating cells on top of the cell layer moving in an independent fashion compared to the cells sheet beneath. Scale bars: 20 μm.

## 4. Discussion

We have in this work investigated the ability of label-free DIC microscopy to study the dynamics of Atlantic salmon keratocyte cells using explant cultures. The collective behavior of keratocyte cells is highly relevant for further understanding of the defense mechanisms of fish skin. Rather than providing conclusive measurements regarding individual cell speed, morphology or chemical constitution, this report aims to highlight the suitability of DIC and label-free microscopy methods for the study both of primary cells and collective cell dynamics.

The collective behavior of cells over different length scales and image resolutions is a challenging target for both manual and automated analysis. The cells are constantly moving, interacting and changing their shape and propagation direction. The cell image contrast is poor compared to the surrounding medium. Although the image contrast can in general be improved using fluorescence imaging techniques, the requirement of fluorescent labeling and the associated increase of phototoxicity alters the biological system and significantly reduces the possible duration of time-lapse studies and the ability to follow a cell population over time.

From our videos, many interesting observations can be made. The large field-of-view time-lapse studies (imaging many adjacent panels every 2 min over several hours), indicate that approaching avalanches in general do not merge, but rather dispersed or avoid each other, even when all cells originated from the same fish. This was observed three out of three times but could be related to the temperature or light exposure during imaging. Observing meeting avalanches was a relatively rare event as it is hard to predict exactly where from a scale an avalanche will emanate and two different scales must be placed at the right distance from one another for observation. We also noted that the progression was very slow – or even completely halted – of the scale explant cultures while kept at 4°C compared to room temperature. This slow keratocyte migration could be a key contributor to the increased presence of skin wounds in farmed fish during the winter season [16].

The single field-of-view videos (imaged every 5 s) were particularly suitable for revealing single-cell dynamics and their interactions. The DIC microscope provided relatively good contrast of even the very thin lamellipodium and long membrane tubes. Cells were observed to turn, rotate, and to move on top of the cell sheets. The microscope’s in-built focus lock system was important to maintain the cells and their thin lamellipodium (extending on the glass substrate) in focus during several hours of imaging.

The videos trigger many questions concerning keratocyte cells and fish biology. Which factors determine the keratocyte path of propagation? Do the cells divide in culture or are they just advancing down from the harvested scales? From where do they derive their energy? Most of our data were acquired of cells in a saline solution only (HBSS with antibiotics), without any nutritional supplements commonly used in cell growth medium and fetal calf serum. Without any growth medium supplements, the cells were observed active for up to 9 days after harvest, contained in only HBSS medium and antibiotics. The cells were kept at 4°C until imaging, then at room temperature. Future studies will investigate for how long after harvest keratocyte cells remain active and further how conditions like temperature or cell culture medium supplements affect the durability of salmon skin cells.

The analytical complexity of label-free bioimage analysis combined with the enormous variability of motion pattern of salmon epithelial cells make it essential to develop suitable tools for automated analysis. This will enable a rigorous quantitative assessment of the cell population effects of e.g. different chemicals, pathogens, nutrients or temperature changes.

In summary, we have shown label-free DIC microscopy as a promising tool to study both single cells and collective dynamics of salmon scale explant cultures. We hope our videos will inspire many future studies of the collective behavior of fish skin cells and the development of analysis software that can accelerate the quantitative assessment of such and similar label-free bioimage videos. The data generated in this study is available on the research data archive DataverseNO [13], and includes both the full image data as well as easy to visualize down-sampled movies to enable a quick overview of the extensive time-lapse data.

## Data availability

Data underlying the results presented in this paper are available from DataverseNO [13].

